# Effects of oral contraceptive pills on brain networks: A replication and extension

**DOI:** 10.1101/2024.10.10.617472

**Authors:** Gino Haase, Jason Liu, Timothy Jordan, Andrea Rapkin, Edythe D. London, Nicole Petersen

**Affiliations:** Department of Clinical Neurosciences, University of Cambridge, Addenbrooke’s Hospital, Cambridge CB2 0SP, United Kingdom; Department of Psychiatry and Biobehavioral Sciences, David Geffen School of Medicine, UCLA, Los Angeles CA; Department of Obstetrics and Gynecology, David Geffen School of Medicine at UCLA, Los Angeles, CA, 90024, USA; Department of Anesthesiology, Emory School of Medicine, Atlanta, GA, USA

**Keywords:** Oral Contraceptives, Functional Connectivity, Birth Control, Replication, seed-based connectivity, dorsolateral prefrontal cortex

## Abstract

Neuroimaging research has identified significant effects of oral contraceptive pills (OCPs) on brain networks. A wide variety of approaches have been employed, largely in observational samples, with few converging results. This study therefore was designed to test for replication and extend this previous work using a randomized, double-blind, placebo-controlled crossover trial of the effects of OCPs on brain networks. Using functional MRI, we focused on brain regions identified in prior studies. Our analyses did not strictly replicate previously reported effects of OCPs on functional connectivity. Exploratory analyses suggested that traditional seed-based approaches may miss broader, network-level effects of OCPs on brain circuits. We applied data-driven, multivariate techniques to assess these network-level changes, A deeper understanding of neural effects of OCPs can be important in helping patients make informed decisions regarding contraception, mitigating unwanted side effects. Such information can also identify potentially confounding effects of OCPs in other neuroimaging investigations.

## 1. Introduction

In the last three decades, functional magnetic resonance imaging (fMRI) has enabled exploration of the neurological effects of commonly used medications (Khalili-Mahani et al., 2017), including those that may not be generally thought of as neurobiologically active. Like other functional imaging techniques, fMRI can map the distribution of drug action, which extends beyond the initial receptor targets of a drug (Jenkins, 2012). Studies on the effects of drugs (and other medical interventions, such as neuromodulation) on the brain have identified neural markers that may underlie the mechanisms by which these interventions elicit certain behaviors (Gerlach et al., 2022). Simultaneously, the rigorous standardization of brain networks (Kong et al., 2024; Uddin et al., 2023) has improved researchers’ ability to evaluate the neural basis of the behavior affected by a certain drug with increasing clarity. The network approach also serves to provide a scale that bridges the knowledge gap between small, regional differences in connectivity and whole-brain activity (Betzel & Bassett, 2017) that correlates to a specific behavior, offering an intermediate approach in determining the effects of pharmaceuticals in eliciting certain behaviors (An et al., 2017; Gudayol-Ferré et al., 2015). Consequently, contemporary imaging research in neuropharmacology is interested in the changes in these networks to understand the interplay between drugs and the behavioral states of the brain. Harnessed effectively, neuroimaging has the potential to refine patient stratification (Etkin et al., 2024), reducing risks of unwanted side effects for patients. Achieving this goal could transform neuroimaging from an exploratory tool into a central driver of precision medicine, bridging the gap between early-stage clinical trials and real-world treatment success.

One group of pharmaceuticals that has been identified as an important target for future research (Beltz, 2024; Petersen et al., 2023) is oral contraceptive pills (OCPs). OCPs are used by approximately 150 million people as of 2020 (United Nations, 2022). While effective in preventing pregnancy, OCPs are also inconsistently associated with self-reported negative behavioral effects (Hall et al., 2012). In some instances, women report emotional lability, irritability, or depressive symptoms (Skovlund et al., 2016, 2018), which can lead to discontinuation within the first few months of use (Rosenberg et al., 1995; Westhoff et al., 2007), whereas other women experience mood benefits (Hamstra et al., 2017; Oinonen & Mazmanian, 2002; Robakis et al., 2019; Schaffir et al., 2016). Understanding the biological and behavioral impacts of OCPs is essential for prospectively identifying women who may experience unwanted adverse effects, allowing for the selection of more tolerable and, consequently, more effective contraceptive options. Furthermore, a thorough understanding of non-contraceptive effects of OCPs underpins accurately communicating these effects to patients, which is essential to informed consent, a fundamental human right.

OCPs, typically comprising synthetic estrogen (ethinyl estradiol) and one of a variety of synthetic progestins, suppress endogenous progesterone and estradiol production. This suppression inhibits the hypothalamic-pituitary axis from secreting luteinizing hormone (LH) and follicle stimulating hormone (FSH), preventing ovulation. Given the widespread use of OCPs, further research on their effects on brain networks is warranted to identify underlying changes in connectivity that cause the behavioral changes experienced by some women who use them. Progress towards identifying such changes may eventually help forewarn populations at risk for premature cessation of OCPs and subsequent unwanted pregnancies.

Currently, there is no behavioral, demographic, or biological parameter that can distinguish between those who will benefit from OCPs and those who experience adverse effects. Brain imaging studies offer a potential solution. Evidence so far shows that OCPs affect brain networks and the structure of brain regions within these networks, however the observed effects vary between studies. OCPs have been associated with increased prefrontal gray matter volume in an observational study (Pletzer et al., 2010), but reduced prefrontal gray matter in a randomized, placebo-controlled trial (Petersen et al., 2021a). Review articles suggest that while OCPs influence brain structure, the direction and location of effects remains unclear. Inconsistent effects of OCPs on resting-state functional connectivity have also been reported (Casto et al., 2022).

A deep phenotyping approach has recently gained traction as a means to more robustly map brain networks (Poldrack et al., 2015), including assessments of hormone-induced changes in brain networks across the menstrual cycle (Pritschet et al., 2020). A recent report found that whole-brain connectivity increases with estradiol levels, and that the dorsal attention network (DAN) and default mode network (DMN) see increased global efficiency at intervals predicted by estradiol levels (Pritschet et al., 2020). Progesterone, on the other hand, had an opposite effect, and largely decreased inter-region connectivity. These results stand somewhat in contrast with a previous deep phenotyping study, which found that progesterone levels were associated with increases in eigenvector centrality, a different but related measurement of connectivity (Arélin et al., 2015). These small but important studies highlight the need for ongoing research to understand the dynamic effects of hormones on brain networks.

Considering regional specificity may identify brain regions that are most influenced by hormone changes. Hence, seed-based resting state functional connectivity (rsFC) analysis is well-suited to investigate brain regions or circuits prone to such volatility. Such studies – investigating the impact of OCPs on seed-based rsFC – are few in number and have produced inconsistent results in identifying regions that displayed significant change (Casto et al., 2022). Three rigorous studies (Engman et al., 2018; Nasseri et al., 2020; Sharma et al., 2020) have directly investigated effects of OCPs on seed-based rsFC.

Nasseri et al. (2020) evaluated rsFC following exposure to a stressor during the active and placebo phases of OCP use, and they found significantly higher left amygdala –ventromedial prefrontal cortex connectivity and lower parahippocampus-lateral occipital cortex connectivity during the hormone-present versus the hormone-absent phase.

Sharma et al. (2020) focused on developmental effects of OCPs, comparing women who initiated use during adolescence vs during adulthood. In both groups of OCP users, connectivity between the right putamen and middle frontal gyrus was higher than in naturally-cycling controls.

Finally, Engman et al. (2018) used a double-blind, placebo-controlled procedure to test the effects of OCs compared to a normal menstrual cycle. The study used a combined oral contraceptive with levonorgestrel, an androgenic progestin, and found a significant decrease in connectivity between the amygdala and the postcentral gyrus in women using OCs compared to the natural cycling and placebo groups. They also found increased connectivity between the dorsal anterior cingulate cortex (dACC) and both the superior frontal gyrus (SFG) and the precuneus when comparing OC usage to the follicular phase of naturally cycling individuals.

In sum, the previous work on seed-based rsFC has implicated numerous regions important to large-scale brain networks necessary for behavioral function. However, the studies evaluated disparate seeds, and results from these studies have yet to be replicated. In this paper, we investigate the effects on functional connectivity between various brain regions during the use of OCPs, considering their role in the context of the larger brain networks they are involved in and the behavioral implications of such networks. The goal of our analysis of this placebo-controlled, double-blind crossover study was to determine whether we could replicate the significant changes in connectivity in the seed regions identified in previous studies to evaluate causal effects of OCPs on brain networks.

Subsequent exploratory analyses were conducted to (1) evaluate relationships between functional networks and subjective symptom reports, and (2) implement a data-driven, multivariate method to test whether more complex models better describe the effects of OCPs on the brain.

To achieve the latter, we applied functional connectome fingerprinting (Finn et al., 2015; Ramduny & Kelly, 2024), a technique that integrates time series data from multiple brain regions to identify stable, individual-specific patterns of connectivity, as well as shared patterns of functional connectivity within a group. This method is particularly well-suited for within-subjects designs, as it reliably identifies individuals across different brain states (e.g., rest and task) and captures connectivity profiles that are not only unique but also predictive of important cognitive traits, such as fluid intelligence. It has also shown some potential as a prognostic tool to predict symptom improvement in major depression (Fan et al., 2020). Therefore, we selected this method to leverage the within-subjects data produced by this randomized, cross-over trial. The goal of this approach was to distinguish features of the connectome unique to individual study participants from those shared across the groups, which could be plausibly attributed to OCP use.

## 2. Methods & Materials

All procedures were approved by the UCLA Medical IRB-3, IRB#16-000396. Written, informed consent was obtained from all participants prior to initiating study procedures

### 2.1 Participants

Detailed recruitment procedures, demographics, and reasons for disqualification at each phase of the study have been previously published (Petersen et al., 2021). Briefly, participants were recruited from the greater Los Angeles community via Craig’s List advertisements. In total, 183 participants (all female) were assessed for eligibility, and of those, 26 ultimately completed both arms of the study. The mean age of the participants who completed the study was 28.4 years (SD = 3.88, range: 20-33 years), and they completed a mean of 15.8 years of education (SD = 1.81, range: 12-21 years). All participants were right-handed according to the Edinburgh Handedness Questionnaire.

Potential participants were excluded if: they were younger than 18 or older than 35; urinalysis or self-report indicated illicit drug use; they self-reported cigarette use; they tested positive or self-reported being currently pregnant or breastfeeding; they had previously used hormonal contraceptives within 3 months of enrolling in the study; they had an ongoing psychiatric disorder according to the Mini International Neuropsychiatric Interview; they reported any major disorder of the central nervous system, or any neurological or endocrine disorders; or they reported any condition that would be contraindicated for safe magnetic resonance imaging. Participants were required to endorse a history of previous negative affect while using OCPs.

### 2.2 Study Design

This study was double-blinded and randomized during data collection. Following the successful completion of an initial screening process, which included a phone-based interview, participants were invited to an in-person session where they received a detailed description of the study design and provided written consent to enroll. Participants were then further assessed for eligibility through a psychiatric evaluation and the completion of drug use and MRI safety questionnaires. Eligible participants were randomly assigned to one of two intervention groups, Group A (OCPs) or Group B (placebo).

Once assigned to an intervention arm, participants were prescribed 21 pills containing either an OCP (30 μg ethinyl estradiol/0.15 mg levonorgestrel) or a visibly identical placebo; both investigational products were prepared by the UCLA research pharmacy. Participants were instructed to begin the intervention on the first day of menses and returned for a testing session after 18-21 days in each intervention arm. Testing sessions included MRI scans, mood-related questionnaires, and hormonal analysis. The latter involved a nurse or phlebotomist drawing approximately 4 mL of blood from participants on testing days, and electrochemiluminescence was used to determine the levels of 17β-estradiol and progesterone, with a detection threshold of 0.03 ng/mL (Roche Elecsys Immunoassay system, F. Hoffman-La Roche, Basel, Switzerland).

Participants then underwent a full menstrual cycle washout period before initiating the opposite intervention arm. The washout period allowed sufficient time for the dissipation of synthetic hormones from the OCP arm, enabling the return of ovulation.

**Figure 1** shows a CONSORT diagram of the study.

**Figure 1.**
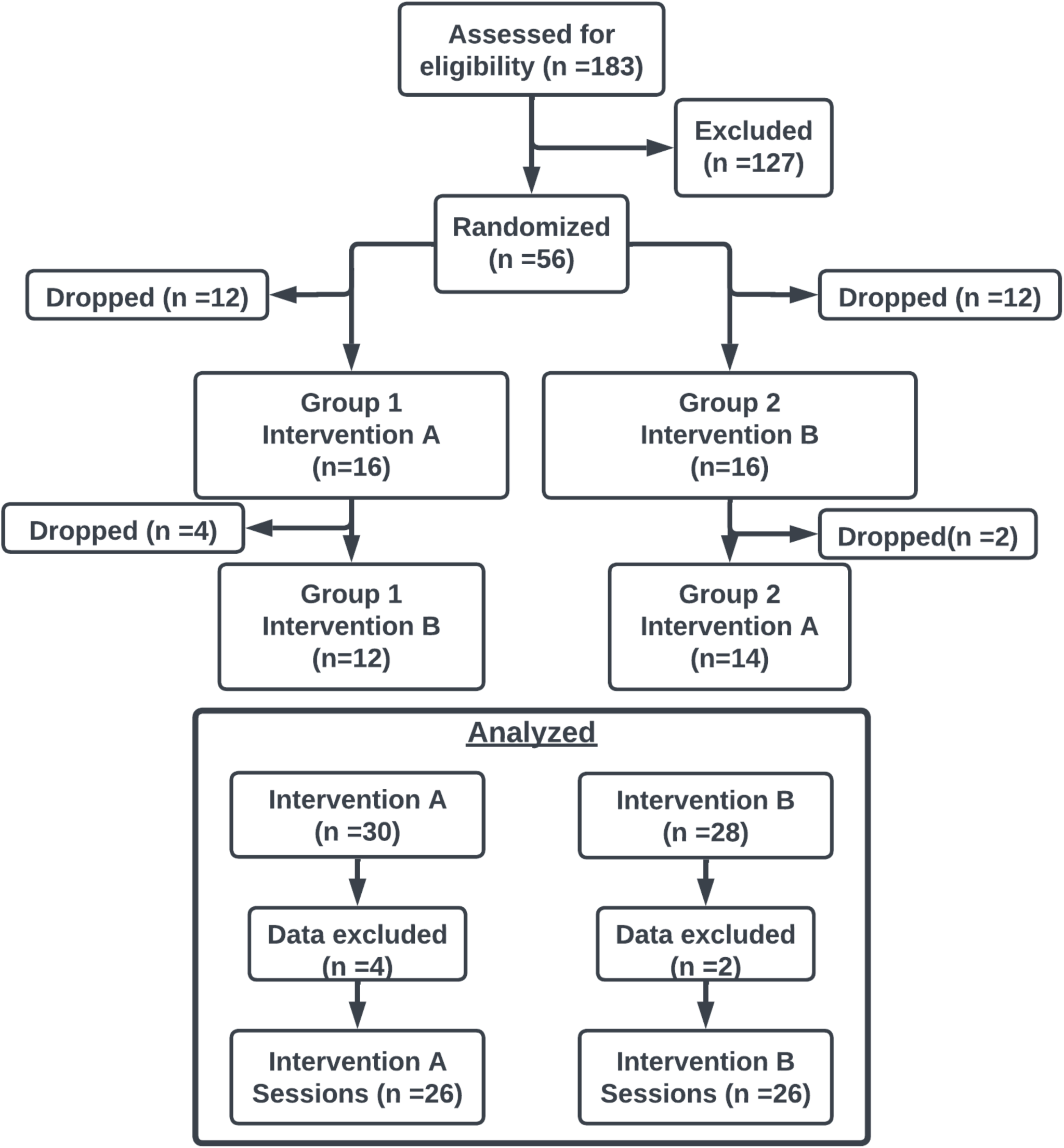
CONSORT diagram depicting overall study flow. Our study design was a randomized double-blind placebo-controlled crossover study. Eligible participants were randomized to either OCPs first and placebo second, or vice versa. Data were included if a participant completed even one session. Three participants were lost to follow-up between the two intervention arms. Data from two participants was excluded due to excess motion during MRI. Brain imaging data from one participant during arm A and one participant during arm B was not collected due to MRI facility time constraints.

### 2.3 Behavioral Data Collection

Questionnaires included the Difficulties in Emotion Regulation Scale (DERS) (Gratz & Roemer, 2004), the Beck Depression Inventory (BDI) (Beck et al., 1988), and the Daily Record of Severity of Problems (DRSP) (Endicott et al., 2006). Participants completed the DRSP online every day of the study, though only the results obtained in the ten days leading up to each MRI scan were included in the analysis, giving time for each intervention to take effect.

### 2.4 Brain Imaging Data Collection & Preprocessing

Whole-brain structural and functional MR imaging was conducted on a 3 Tesla Siemens Prisma Fit MRI scanner with a 32-channel head coil at the UCLA Staglin Center for Cognitive Neuroscience. A single T1-weighted structural scan (TE= 2.24ms; TR= 2400ms; voxel resolution= 0.8 x 0.8 x 0.8 mm^3^) was collected during the intake session as well as a 8 minute baseline T2*-weighted multi-band sequence resting state functional scan. Resting state functional scans (TE= 37ms; TR= 800ms; FoV = 208mm; Slice Thickness= 2mm; Number of Slices = 72, voxel resolution= 2 x 2 x 2 mm^3^) were performed on test days (18-21 days after beginning OCPs or Placebo). Prior to all resting state functional scans, two spin echo fieldmaps were collected in opposite directions (AP and PA).

Processing of fMRI data was carried out using FEAT (FMRI Expert Analysis Tool) Version 6.00, part of FSL (FMRIB’s Software Library, www.fmrib.ox.ac.uk/fsl). The following pre-statistics processing was applied; motion correction using MCFLIRT(Jenkinson et al., 2002); B0 unwarping using boundary-based registration via FUGUE (Jenkinson, 2003, 2004); slice-timing correction using Fourier-space time-series phase-shifting; non-brain removal using BET (Smith, 2002); spatial smoothing using a Gaussian kernel of FWHM 4.0mm; grand-mean intensity normalization of the entire 4D dataset by a single multiplicative factor; highpass temporal filtering (Gaussian-weighted least-squares straight line fitting, with sigma=50.0s). ICA-based exploratory data analysis was carried out using MELODIC (Beckmann & Smith, 2004), in order to investigate the possible presence of unexpected artifacts or activation. ICA-FIX was trained on a set of 20 scans that were hand-classified into noise and non-noise components, with the scans randomly selected from 5 bins sorting scans by the amount of average motion present to have high and low motion data in the trained set. The component classification derived from the trained data was then used in ICA-FIX to classify noise and non-noise components from all subject data and non-aggressively remove the noise components. After denoising, ICA-FIX applied a high-pass filter to each subject’s data. Registration to high resolution structural and/or standard space images was carried out using FLIRT (Jenkinson et al., 2002; Jenkinson & Smith, 2001). Registration from high resolution structural to standard space was then further refined using FNIRT nonlinear registration (Andersson et al., 2007b, 2007a). Lastly, average time series were extracted from each subject’s data based on brain nodes specified by the atlas then detrended for cubic trends and finally z-score normalized.

### 2.5 Seed info

We used the brain parcellation proposed by Van De Ville (Van De Ville et al., 2021) to extract time series from all imaging data. Briefly, this parcellation includes a Schaefer 400 brain region cortical parcellation ((Schaefer et al., 2018), https://github.com/ThomasYeoLab/CBIG/tree/master/stable_projects/brain_parcellation/Schaefer2018_LocalGlobal) combined with 16 subcortical regions and 3 cerebellar regions from the HCP release for a total of 419 nodes.

We selected four seeds from three previous studies to attempt to replicate previous reports of OCP effects on functional connectivity in these regions: the left amygdala (Nasseri et al., 2020; Engman et al., 2019), right putamen (Sharma et al., 2020), left parahippocampal gyrus (Engman et al., 2019), and dorsal anterior cingulate cortex (dACC; Engman et al., 2019) as seeds. We determined that these corresponded approximately to nodes 406 (amygdala), 411 (putamen), 405 (parahippocampal gyrus), and a combination of 360+177 (dACC). These nodes are shown in Figure 2.

**Figure 2:**
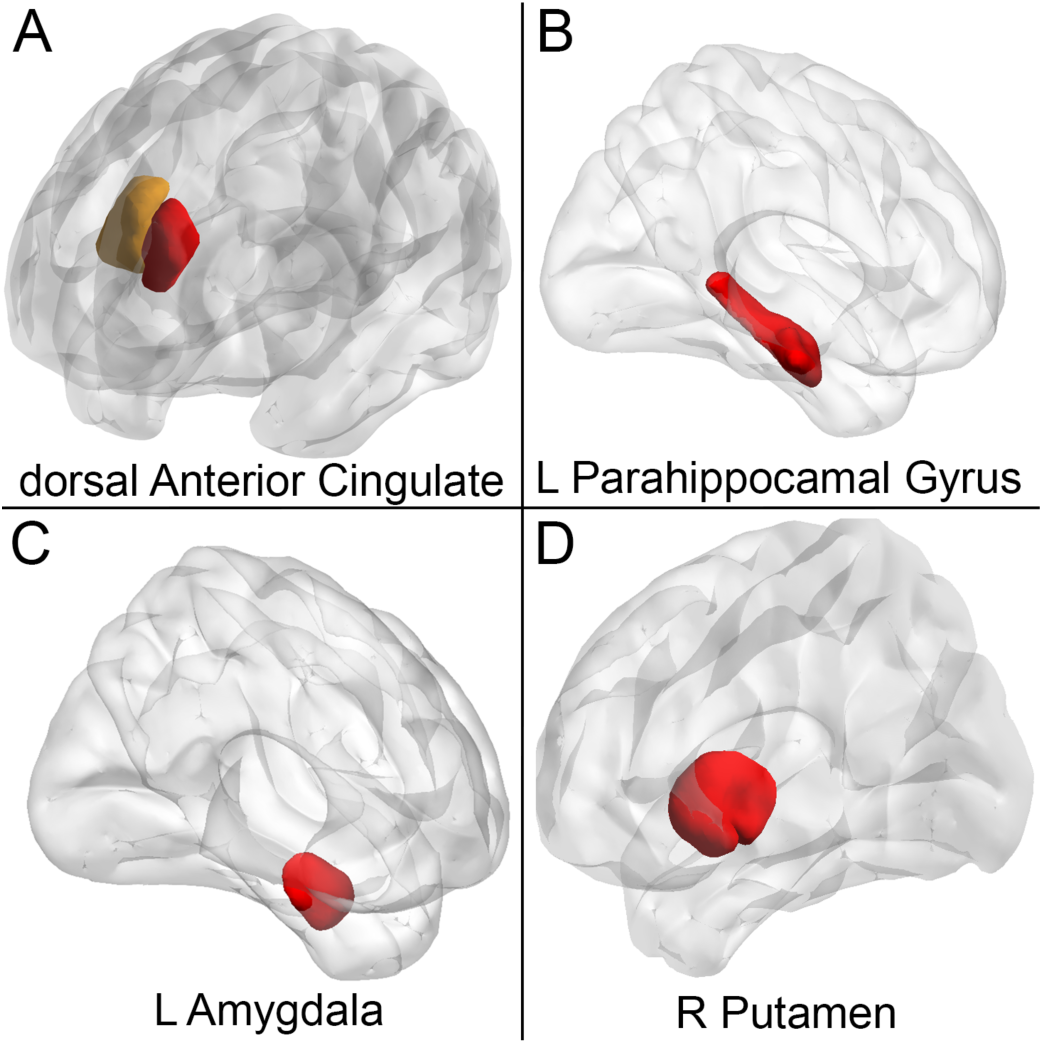
Regions of interest (ROIs) used for seed-based resting-state functional connectivity analysis, displayed on a standard brain template. Seeds were selected based on prior research and include the left amygdala (node 406), right putamen (node 411), left parahippocampal gyrus (node 405), and dorsal anterior cingulate cortex (dACC; combination of nodes 360+177). These seeds were derived from the Schaefer atlas combined with subcortical nodes as previously described to assess the effects of oral contraceptive pills (OCPs) on functional connectivity.

### 2.7 Analyses

#### 2.7.1 Functional Connectivity Analysis

For each participant, each session, seed-based functional connectivity analysis was performed using FMRIB Software Library (FSL). Following data preprocessing and cleaning as described above, mean time series were extracted from each ROI and used as the seed time series. Correlations between the seed time series and all other voxels in the brain were calculated to determine connectivity.

To evaluate the effect of OCPs on functional connectivity of *a priori* seeds, mean time series were extracted from each relevant node in the atlas and Pearson’s correlation between each node-to-node pair was calculated. Functional connectivity values were converted to standardized z scores using a Fischer transformation and then compared between sessions (OCP vs. placebo) using a one sample t-test.

Because the parcellation scheme employed also generates edges reflecting connectivity between every other node-node pair in the brain, we also calculated the effect size of each change (OCP to placebo) using Cohen’s *d.* We averaged together Cohen’s *d* values for each node in 7 canonical networks as previously defined (Thomas Yeo et al., 2011).

#### 2.7.2 Functional Connectome Fingerprinting

Functional connectome fingerprinting was used to compare changes in identifiability and idiosyncrasy across the whole brain connectome between the two study arms. **Identifiability** refers to the capacity to distinguish individual-specific patterns of brain connectivity from group-level data, where each participant’s brain network profile remains consistent across different scans or conditions. **Idiosyncrasy** refers to the degree to which specific brain connectivity patterns are unique to an individual compared to others in the group. High idiosyncrasy means that the brain connectivity profile of an individual is more distinct and less likely to resemble those of other participants.

Functional connectome fingerprinting integrates these concepts, and allows investigators to track how stable or unique a participant’s brain connectivity is between different states, such as during OCP and placebo phases. This technique leverages individual differences in neural network activity to investigate the level of consistency of functional connectivity edges, or connections between nodes, in the functional connectivity data collected from one subject session as compared to a separate retest, intratest, or group-level dataset (Amico & Goñi, 2018; Finn et al., 2015). Nodes represent specific brain regions, while edges represent the functional connectivity (i.e., correlation of activity) between these regions. Amico et. al expanded this test-retest fingerprint to work within a single session by comparing the first and second half of the scan. This approach allows determination of idiosyncrasy and identifiability to be determined without multiple sessions. When this technique is used for comparison of within-scan examination for the same subjects under two different conditions, it allows for determination of how the condition alters identifiability and idiosyncrasy in the respective group. Using the spatial (Pearson) correlations between each half-scan (first 4 minutes and last 4 minutes of the 8 minute scan sequences for both OCP and placebo arms), we were able to determine each subject’s self identification *I_Self_*, identification with others *I_Other_*, and their overall identifiability *I_Diff_* according to each condition (OCP half 1, OCP half 2, placebo half 1, placebo half 2). These are defined as follows:

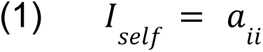

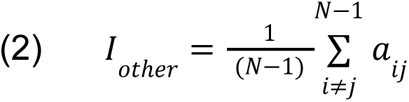

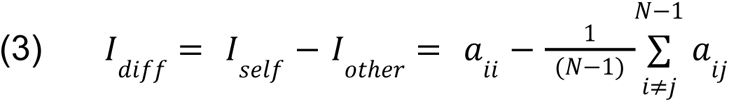

These three metrics measure the correlation between a target half-scan and the other half scans in the identifiability matrix **A. Matrix A** has the dimension **N^2^**, where **N** is the number of participants. We can then denote **a_ii_**as a measure for the correlation between half-scans of the same subject, effectively including only the diagonal elements of **A**. We can use this as a measure of self-identifiability, making *I_Self_* equal to the value **a_ii_** for each subject. Similarly, *I_Other_* is the average of **a_ij_**, which contains all off-diagonal, or non-self, correlations; *I_Diff_* is the difference between *I_Self_* and *I_Other._* The use of the half-scan method thus makes it possible to obtain each subject’s metrics between OCP and placebo without the need for multiple scans. Further, using intraclass correlations (ICC) (Bartko, 1966), we are able to determine which connections drive this identifiability. Such connections that have high ICC values are considered idiosyncratic connections, since their value contributes disproportionately highly to the ability to identify a subject’s connectome among a group. The code for these calculations was previously made available by Tolle et al. (https://github.com/eamico/Psilocybin_fingerprints) and our implementation is stored in our above-named repository.

#### 2.7.3 Correlating brain and behavior changes

Given the very large size of the dataset (419 x 419 nodes, or 175,561 features per participant), we sought to reduce the total number of features entered into our exploratory search for brain-behavior relationships. Therefore, we began by thresholding the group-level connectivity (i.e., the matrix containing the change in connectivity between each edge) at *p* < 0.001. Next, we entered these remaining features into a principal component analysis using the PCA command in the python library *scikit-learn*. Because of this conservative thresholding, the only features entered into the principal component analysis were those that changed substantially as a result of OCP use. Subsequently, we correlated the first principal component with the change in DRSP scores for each participant. Finally, we correlated each individual feature (edge) included in the first principal component against each participant’s DRSP scores using Pearson correlations.

## 3. Results

### 3.1 Replication Findings

In replicating the methods of the three aforementioned studies (Engman et al., 2018; Nasseri et al., 2020; Sharma et al., 2020), we analyzed the connectivity differences between the OCP and placebo arms using Schaefer nodes associated with each respective ROI seed in a nodewise seed-based whole-brain comparison. Visualizations of each ROI node are shown above in **Figure 2**. For completeness, we carried out the same analysis with a different ROI definition, shown in **Supplemental Figure 1**, obtaining essentially the same overall results.

To test for replication of the results of Nasseri et al., we identified the Schaefer nodes corresponding to the left amygdala and parahippocampal gyrus seed region masks that were used in their analysis. We ran a seed-based connectivity analysis of the whole brain based on those nodes, focusing on ROI-ROI connections that were previously reported to differ significantly in OCP users. We found that, compared to placebo, during active OCP treatment, the left amygdala had no notable increase in connectivity to the ventromedial prefrontal cortex (*p* = 0.741; *t* = –0.335). Additionally, the left parahippocampal gyrus did not show a decrease in connectivity to the right superior lateral occipital cortex defined as Schaefer nodes 285 or 282 (*p*s > 0.05). Expanding our search beyond these (previously-investigated) ROIs to all other nodes in our parcellation, we found no changes in left amygdala or parahippocampal gyrus connectivity between the two study arms, even at an uncorrected threshold.

Next, we used the time series from the subcortical node corresponding to the right putamen seed to test for replication of the results of Sharma et al. At an uncorrected threshold, we found a significant increase in connectivity to a node in the middle frontal gyrus (Schaefer node 186; *p* = 0.0481; *t* = 2.18), which is consistent with the findings of Sharma et al., however the node we found was contralateral to their resulting region (visualized in **Supplemental Figure S2**). The connectivity between the right putamen and the middle frontal gyrus node corresponding to the MNI coordinates reported by Sharma et al. (30, 12, 60) did not show any significant changes in connectivity (*p* = 0.586; *t* = 0.553).

Finally, we used the time series data extracted from the Schaefer nodes corresponding to the left and right dACC regions to replicate the results from the study by Engman et al. (2018). We did not replicate the reported increase in connectivity to the superior frontal gyrus (*p* = 0.846; *t* = 0.197) from the right dACC, though at an uncorrected threshold, we did find a decrease in connectivity from the right dACC to the adjacent right frontal medial cortex (*p* = 0.0419; *t* = –2.17). Further, the connectivity of the left dACC node did not replicate the lower connectivity to the precuneus location (MNI: 0, –54, 63) found by Engman et al. (*p* = 0.718; *t* = 0.367), though we did find a decrease in connectivity to a nearby right precentral gyrus node (*p* = 0.0264; *t* = –2.39; Schaefer node 195; comparison shown in **Supplemental Figure S3**) from the placebo to the OCP arm. Additionally, as reported above, we were not able to replicate changes in amygdala connectivity (no significant change in the connectivity to the right postcentral gyrus or nearby nodes, *p* = 0.419; *t* = –0.825).

### 3.1 Exploratory Findings 1: Alternative connectivity changes

In the absence of replicating previously-reported effects, we did a data-driven exploration to look for other effects of OCPs on functional connectivity. This exploratory analysis can be considered hypothesis-generating rather than hypothesis-confirming.

Effect sizes were calculated to describe the difference in connectivity between each Schaefer node and each other Schaefer node as a result of OCP. Each node was used as a seed and these seeds were grouped into their respective brain networks according to the division by Yeo et al (7 network division). Figure 3 presents a matrix of Cohen’s *D* effect sizes from this analysis. The highest frequency of medium to large effect sizes occurred in subcortical networks, along with executive network and somatomotor network nodes. Each node-node connectivity change is depicted in Figure 3, panel A, and a summary visualization showing the average network-network connectivity change is shown in panel B. Positive (red) average Cohen’s *D* values indicate increased connectivity in the OCP arm compared to placebo. Negative (blue) values indicate network changes that had a decrease from the placebo to OCP arms.

**Figure 3:**
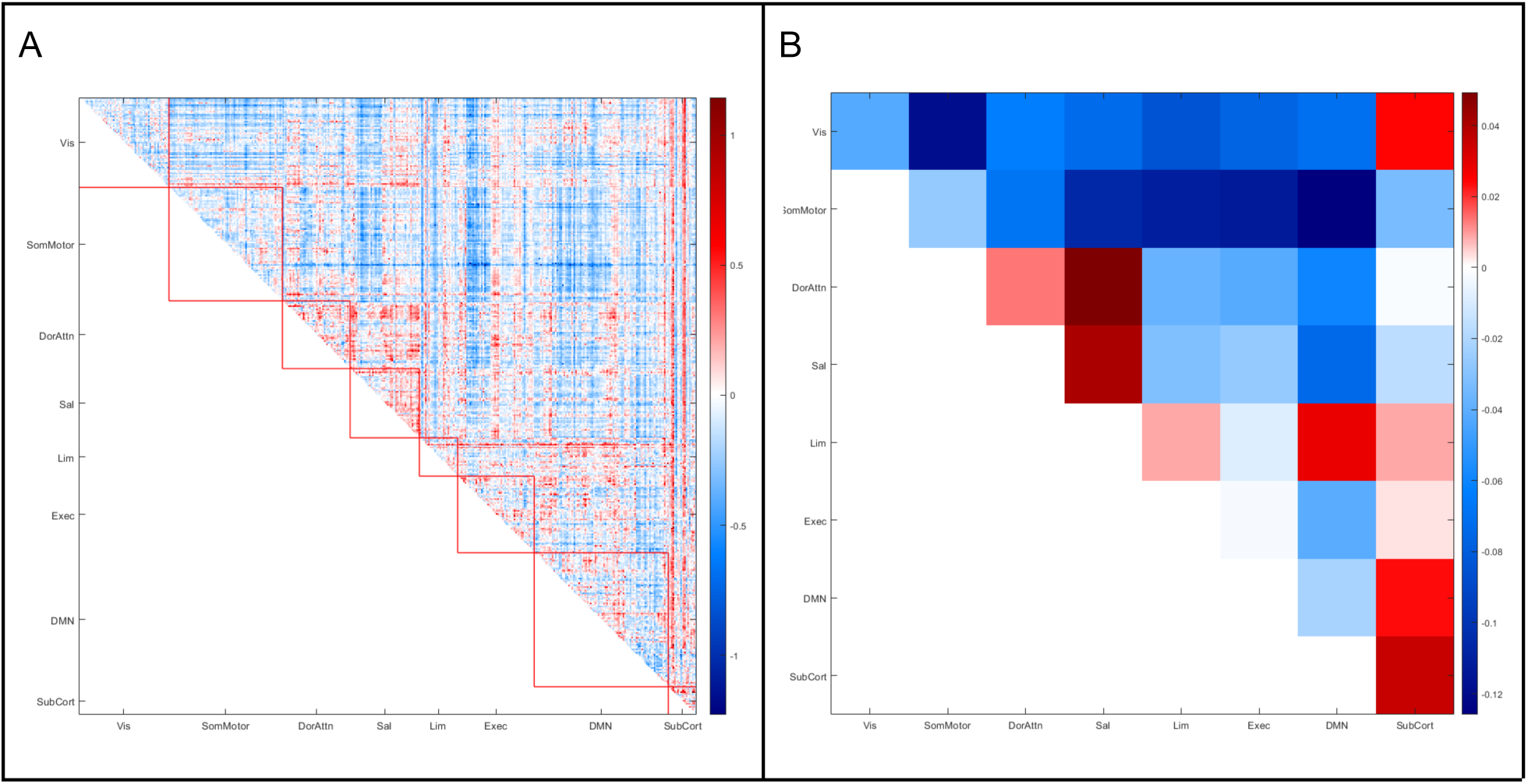
Effect size (Cohen’s d) matrix depicting connectivity changes between Schaefer atlas nodes during the OCP and placebo interventions. The matrix visualizes network-level changes, with increased connectivity (red) and decreased connectivity (blue) identified between each brain node as a result of OCP use. Notably, the subcortical, executive, and somatomotor networks exhibited the largest effects. Panel B summarizes the average connectivity change for each network.

### 3.2 Exploratory Findings 2: The functional connectome fingerprint of OCP use

We examined the functional connectome fingerprint of each participant during each intervention to see if OCPs altered identifiability. We found that *I_Other_* was significantly higher (Bonferroni-corrected *p* < 0.001) for participants during the OCP intervention, while *I_Self_* and *I_Diff_* did not differ significantly between intervention arms. This indicates that while OCPs cause participants’ connectomes to become more similar to each other, each individual can still be accurately identified compared to others regardless of intervention. **Figure 4** shows the changes in identifiability between interventions and **Figure 5** shows an identifiability matrix. This identifiability matrix shows the identifiability of each half-scan of every participant compared to the subsequent half-scans. Participants are placed in the same order in each quadrant’s axes. The diagonals of every quadrant show self-identifiability of the subject between scan halves. Each row and each column correspond to a participant. Each participant is represented four times in two rows and two columns, which in turn correspond to the two sessions and the two halves of each session for a total of 4 representations within the matrix. Yellow indicates higher correlation between two connectomes, and blue indicates lower correlation between two connectomes. As anticipated, most individuals show high correlations between the first and second half of the same scan, indicated by the yellow diagonal. However, this diagonal dissipates when comparing halves of scans collected during the different study arms. matrix showing identifiability of each half-scan of every subject compared to other half-scans. Subjects are placed in the same order in each quadrant’s axes, so diagonals of every quadrant show self-identifiability of the subject between scan halves.

**Figure 4.**
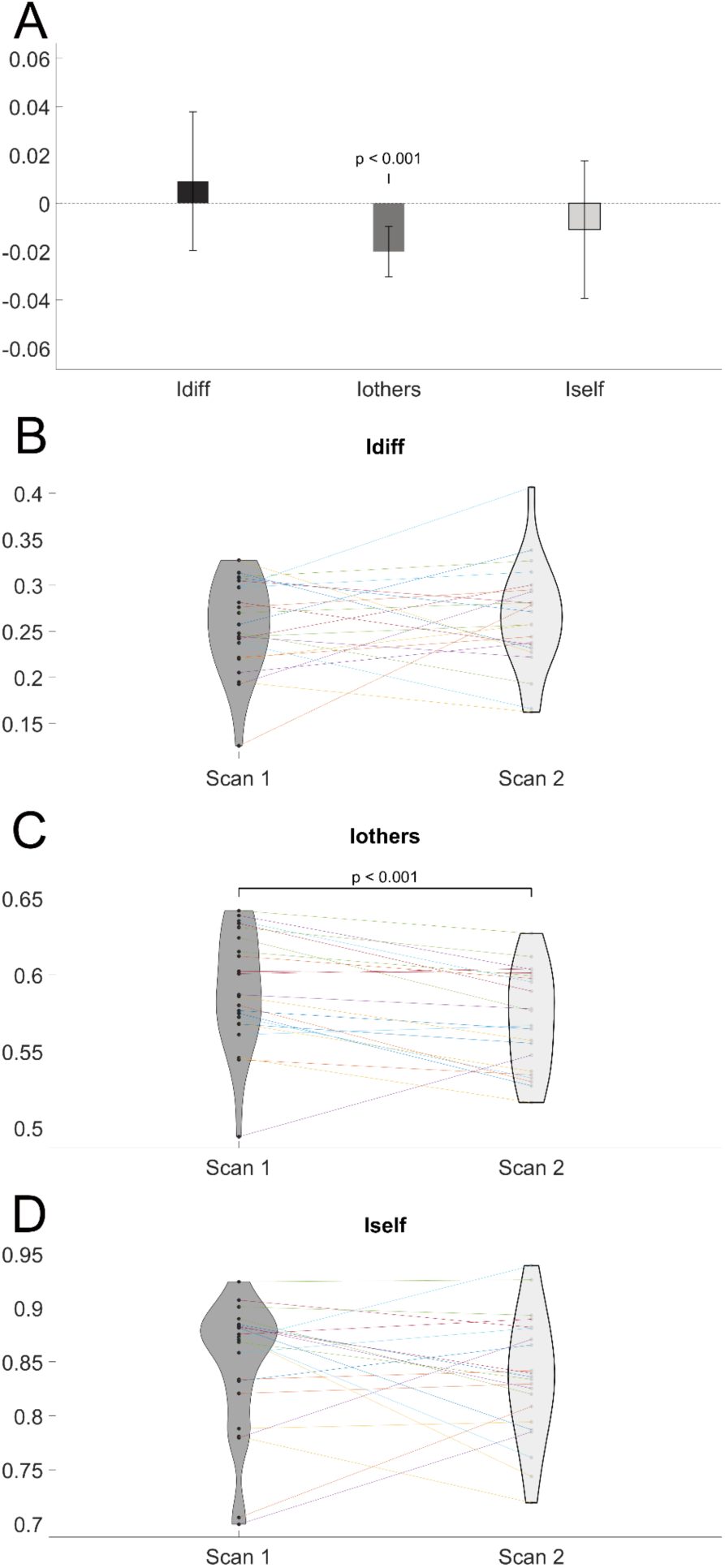
Changes in *I_Diff_*, *I_Other_*, and *I_Self_* between OCP intervention arm (labeled Scan 1) and placebo arm (labeled Scan 2). *I_Self_* measures within-subject consistency, while *I_Other_*reflects between-subject similarity. The results show a significant change in *I_Other_* during OCP use, indicating greater similarity in brain connectivity profiles across participants while taking OCPs.

**Figure 5.**
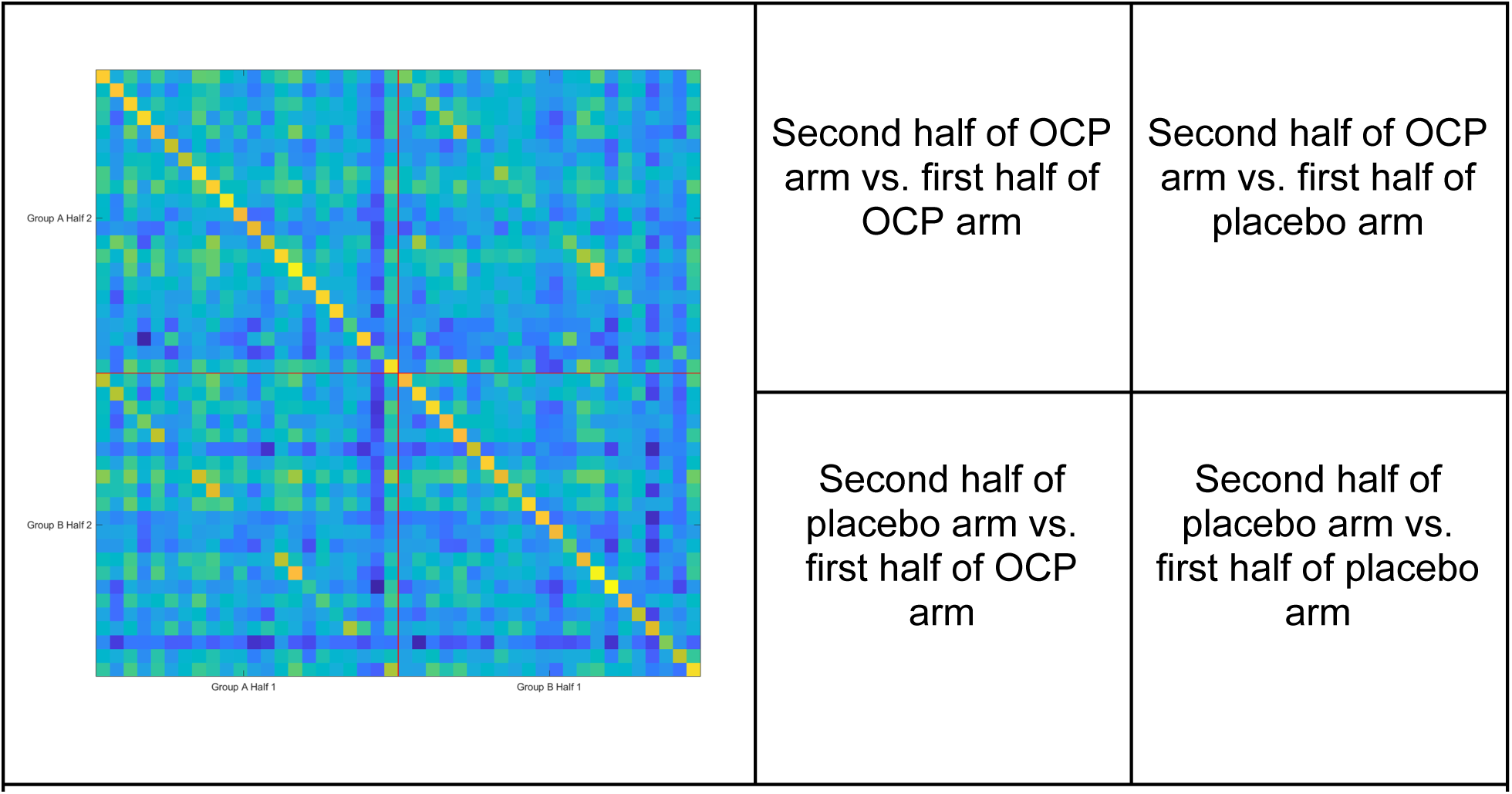
Identifiability matrix showing the correlation between each participant’s brain connectivity profiles across scan halves. Each subject is represented by four entries: two from the OCP arm and two from the placebo arm. The yellow diagonal indicates high within-subject identifiability (self-identifiability), which dissipates when comparing halves of scans between OCP and placebo interventions, highlighting the influence of OCPs on network-level idiosyncrasies.

To understand which brain circuits drove these connectome changes – i.e., examining the idiosyncratic changes due to OCP use – we found significant differences in ICC values for 26 network-to-network connections. Four network-network ICC comparisons were significantly higher (Bonferroni corrected-p < 0.001) for the OCP intervention compared to placebo. All other network-network ICC comparisons were found to be significantly higher (Bonferroni corrected-p < 0.001) for placebo compared to OCP. **Figure 6** shows a heatmap of network-network ICC differences with red indicating higher for OCP and blue indicating higher for placebo. Examining the ICC strength of each network, which is done by summing each network’s total node ICC value from all connections, we found that idiosyncrasy was significantly higher (Bonferroni corrected-p < 0.05) for placebo compared to OCP for all networks except for the visual and dorsal attention networks, as shown in **Figure 7**.

**Figure 6.**
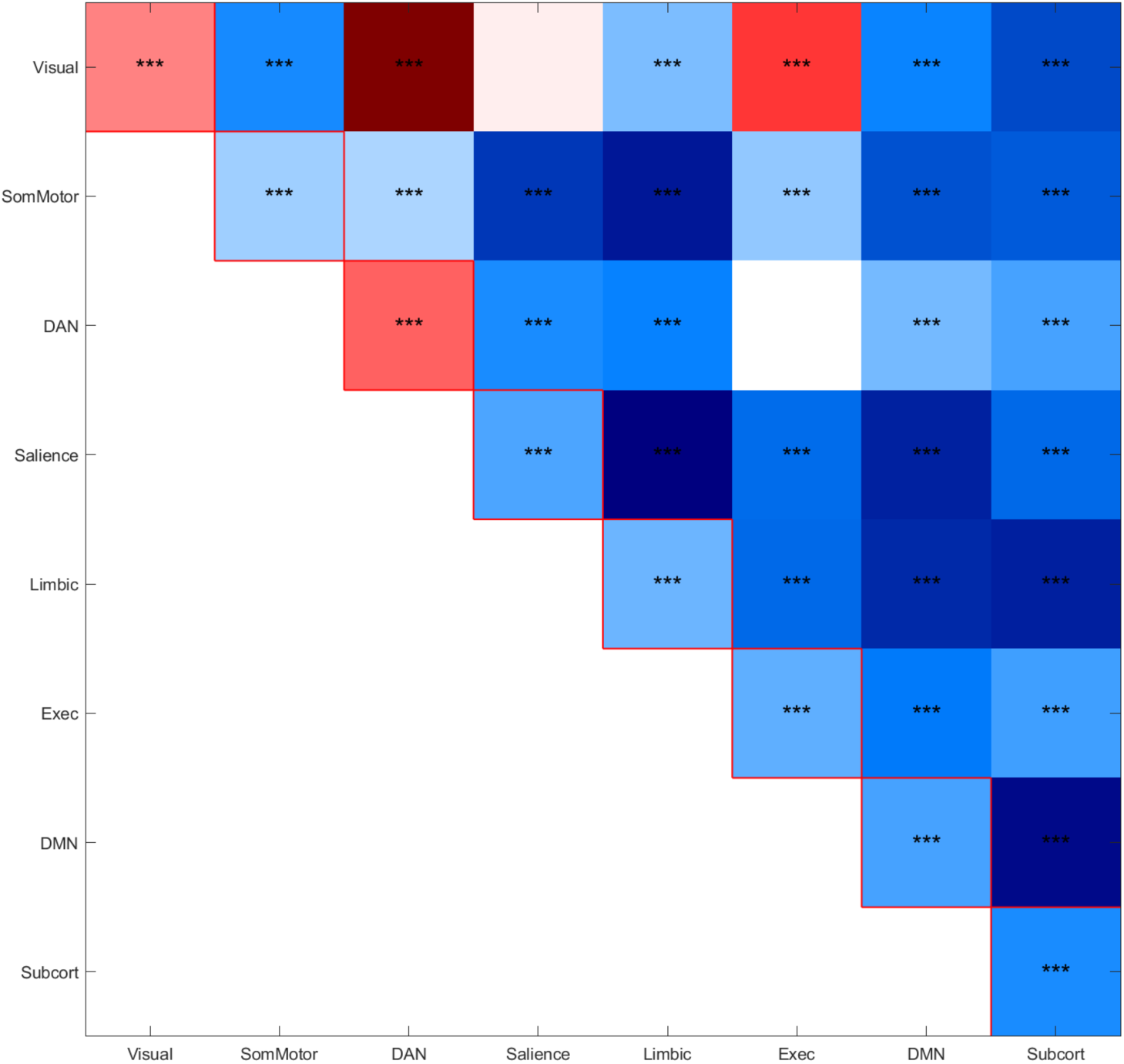
Heatmap showing network-to-network intraclass correlation coefficient (ICC) differences between the OCP and placebo arms. Red regions indicate higher ICC values during OCP use, while blue regions indicate higher values during the placebo phase. Nearly all networks exhibited significant differences in idiosyncrasy, with OCP use reducing the stability of network connectivity patterns.

**Figure 7.**
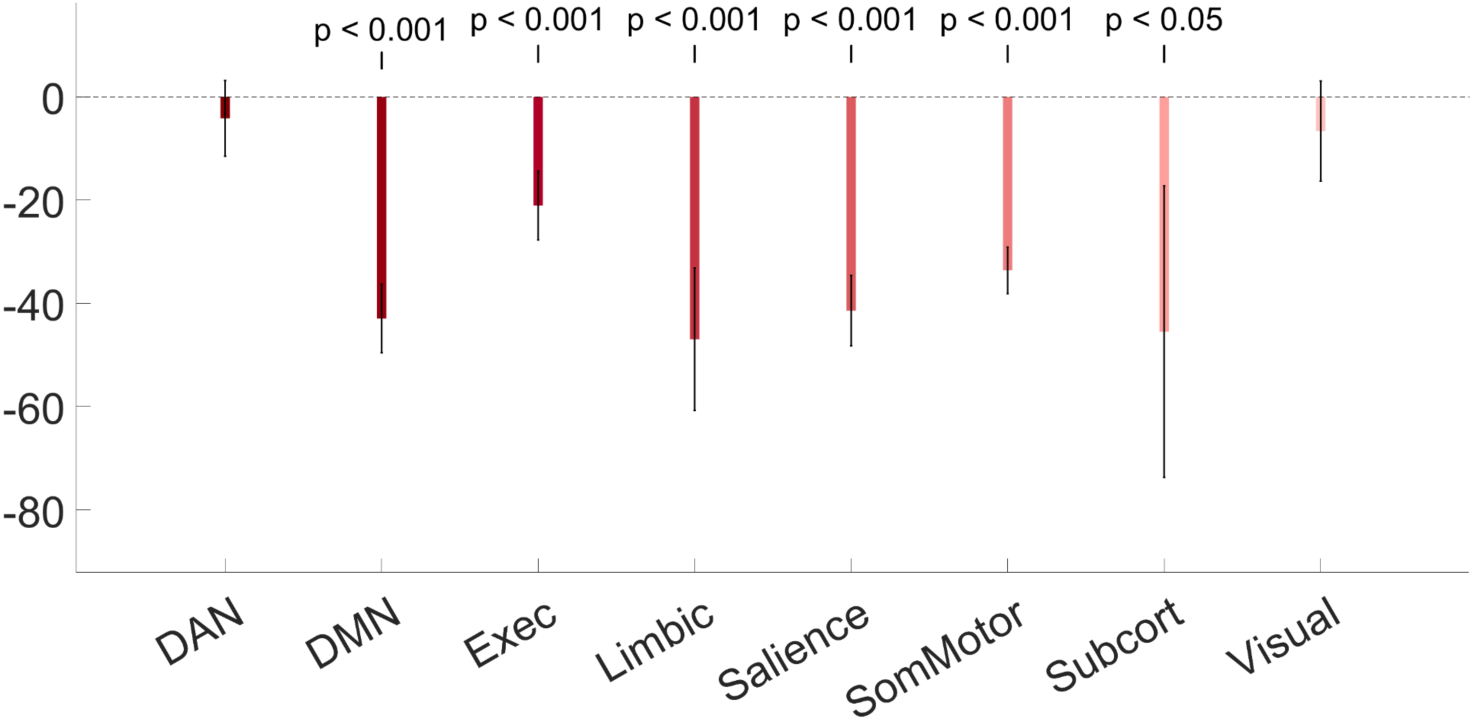
Summed ICC strength for each network comparing the OCP and placebo arms. Overall ICC strength was significantly reduced during OCP use in six out of eight networks, including the somatomotor and default mode networks. The visual and dorsal attention networks showed minimal differences between interventions.

Finally, we evaluated the contributions of idiosyncratic connections to each fingerprint. We achieved this by calculating the identifiability measures using the connections with the top ICC values (i.e., the most idiosyncratic) starting from the top 100 connections until we included the whole connectome again. We observed that until approximately 90% of the connectome is included, the OCP intervention makes participants much less identifiable than when taking placebo. This observation suggests that the most idiosyncratic connections are also the most susceptible to the influence of OCPs.

### 3.3 : Exploratory findings 3: Relating functional connectivity changes to changes in negative affect

In the same dataset, we previously reported (Petersen et al., 2021) that OCPs increased scores on the Daily Record of Severity of Problems (DRSP), with effect sizes ranging from *D* = 0.16 (DRSP item 7, Fatigue, smallest effect) to *D* = 0.88, DRSP item 11, Physical Symptoms, largest effect). This previous analysis did not reveal correlations between changes to brain structures and DRSP items. Therefore, here we explored the possibility that changes to brain *network activity* rather than individual structures better explained the observed increases in DRSP scores.

Given the very large feature set available, we performed data reduction in two steps. First, features were thresholded (arbitrarily) at p < 0.001 to include only edges that changed substantially in response to OCPs. Subsequently, when PCA was applied to this feature set, we found that (regardless of the number of PCs specified), only PC1 correlated with change in DRSP scores, *r* = –0.756, *p* = 7.811e-06. However, the feature weights were overall quite similar to one another. The cumulative loadings are shown in Figure 9.

**Figure 8:**
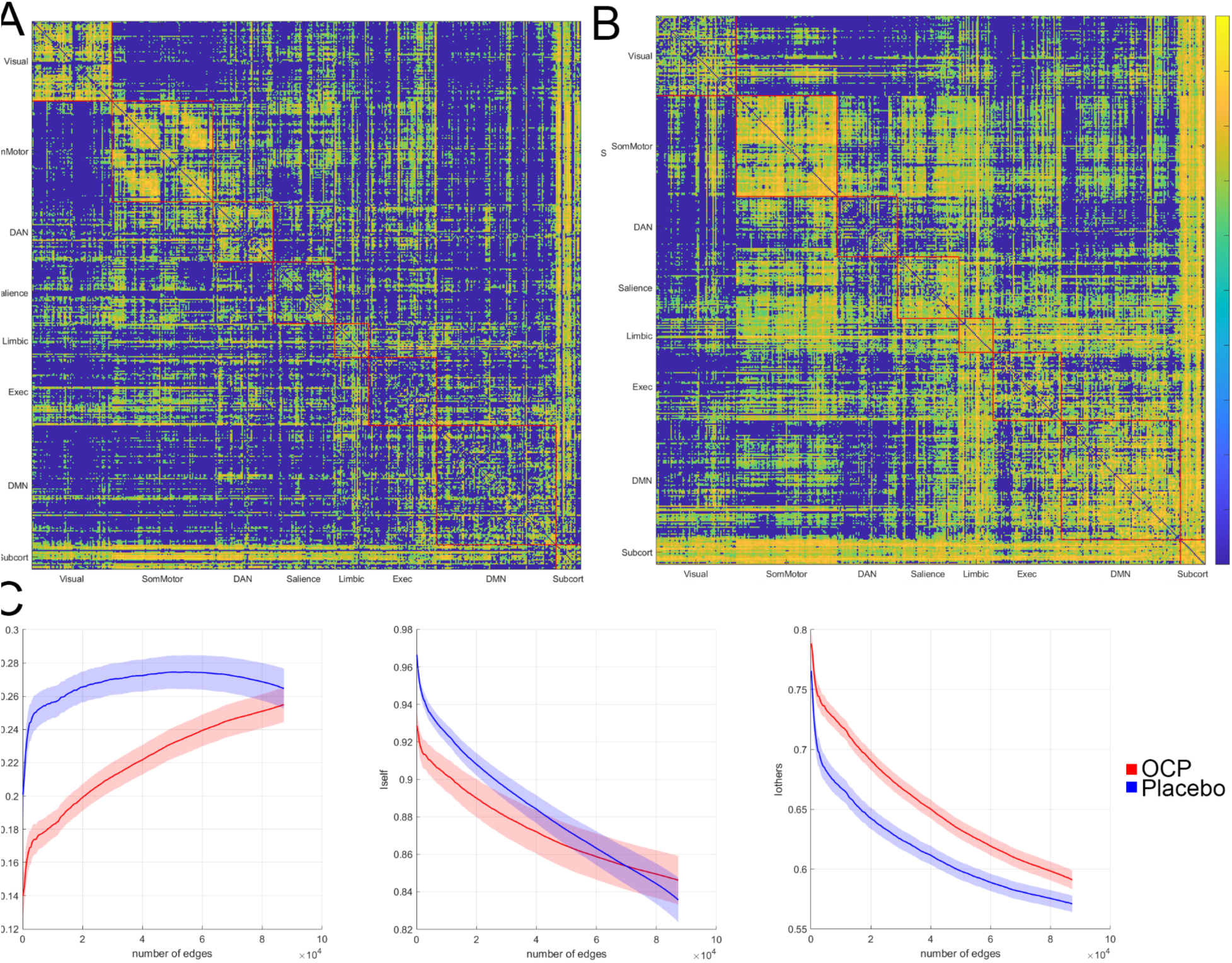
Idiosyncratic contributions to the functional connectome on OCPs and placebo. *A)* ICC matrix of node connectivity measured during OCP intervention, grouped by network. Yellow indicates higher ICC values – i.e., connections that are stable from the first half of an individual’s scan to the second half, but are unique *between* individuals. *B)* Similar ICC matrix of connectivity measured during placebo arm. *C)* Identifiability measured as a function of progressive addition of edges. Edges are added in the order of their ICC value for the placebo intervention in order to examine the effects of OCPs on each subject’s most idiosyncratic connections.

**Figure 9:**
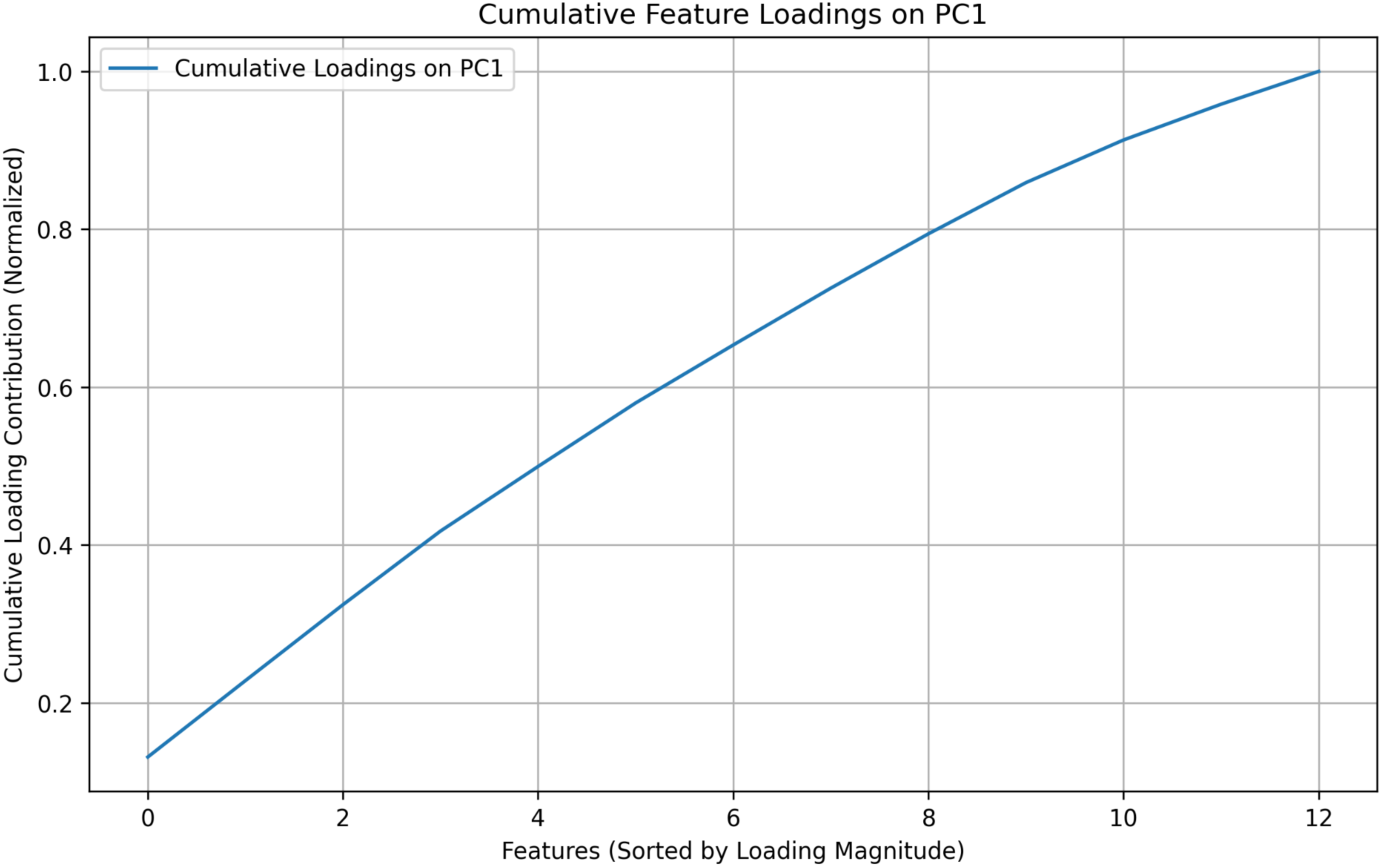
Cumulative feature loadings of a principal component associated with DRSP scores. Network features that changed in response to OCP use at a p < 0.001 threshold were entered into a principal component analysis. The cumulative feature loadings on the first principal component (which explained the majority of the variance, and correlated with DRSP scores) are shown here. The smooth transitions between features and similar slopes indicate that each feature imposes a similar loading on this component.

Examining these features, we found that 13 features (edges) survived the *p* < 0.001 threshold. We plotted the change in each node-node connection against the change in DRSP scores, and observed remarkably similar correlation maps, as shown in Figure 10. Here, each correlation is overlaid on top of each other. Individual plots can be viewed in Supplemental Materials (Supplemental Figure S5).

**Figure 10:**
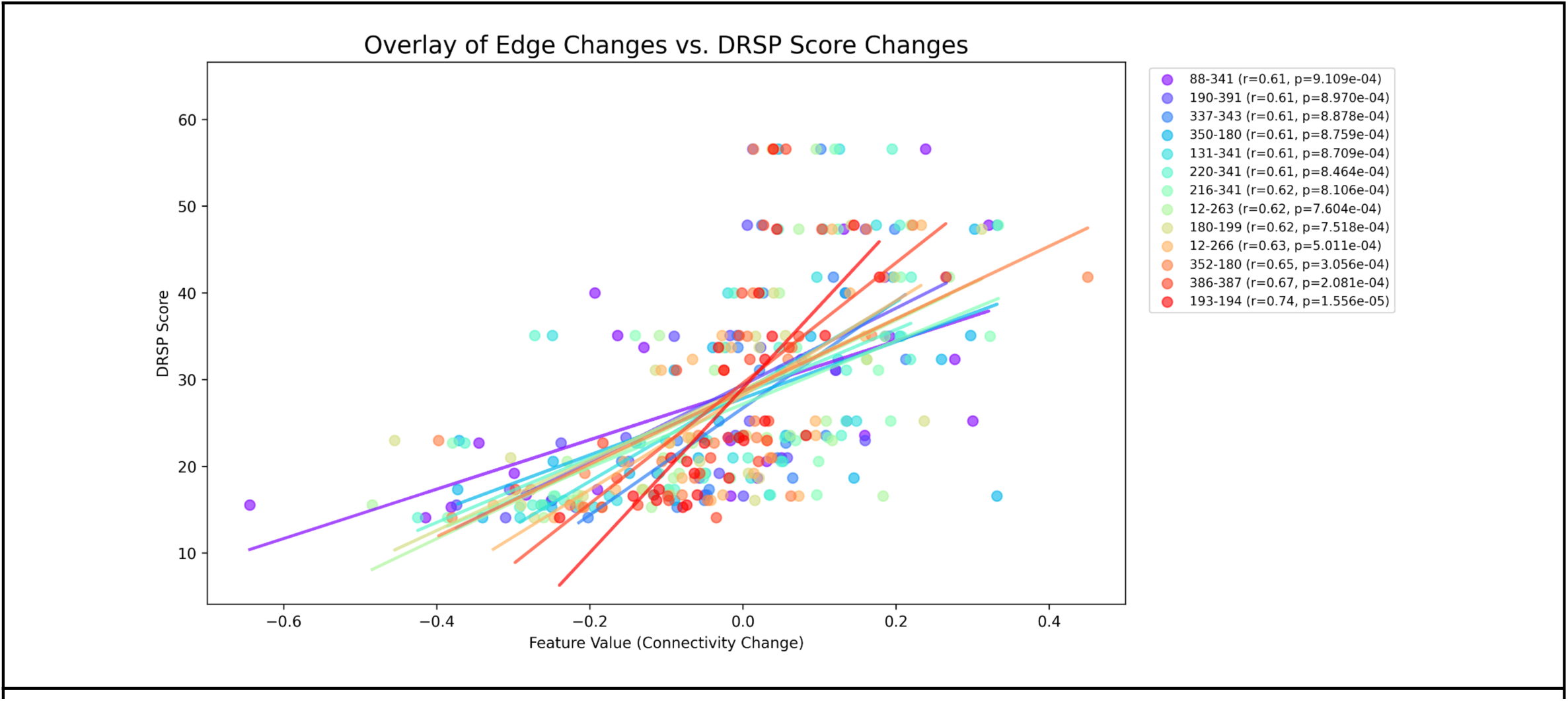
Scatterplot illustrating the correlation between changes in functional connectivity (node-to-node connections) and changes in Daily Record of Severity of Problems (DRSP) scores for 13 features (edges). Each colored line represents a different correlation between edge changes and DRSP changes, with the corresponding Pearson correlation coefficient (r) and p-values shown in the legend. Higher DRSP scores were associated with larger changes in connectivity between the nodes indicated here, suggesting that these connectivity changes may underlie the negative affective symptoms reported during the OCP arm of the study.

The brain regions involved in these correlations include the frontal pole (appears bilaterally in 8 of 13 correlations), right superior frontal gyrus (in 3 of 13 correlations), right middle frontal gyrus (in 2 of 13 correlations), left precuneus (in 2 of 13 correlations), left posterior cingulate (in 2 of 13 correlations), left angular gyrus, and regions of occipital cortex. We suggest that these regions may form a hormone-sensitive network that underlies OCP effects on mood, shown in Figure 11.

## 4. Discussion & Conclusion

### 4.1 Replication Findings

The primary goal of this study was to investigate the replicability of previous findings on the effects of OCPs on seed-based functional connectivity. None of the seeds chosen *a priori* showed changes from the OCP active arm to the placebo arm in the same direction and location as previously reported. Given the diversity of study designs, OCP formulations, and seed selection approaches in prior work, it is unsurprising that our replication efforts yielded mixed results. Below, we compare our findings to three key studies, highlighting both consistencies and discrepancies, and suggest possible explanations based on methodological differences. We also report exploratory findings showing widespread influences of OCPs on functional connectomes. We interpret these data as evidence that OCPs produce diffuse, rather than focal, influences on brain networks. We offer potential regions for inclusion in such analyses in future explicit replication studies.

Using a strict interpretation, our explicit replication was not able to verify previous reports of seed-based connectivity differences between women who did and did not use active OCPs. However, there were some notable similarities in our results, including an increase in putamen-*left* middle frontal gyrus connectivity (vs. previously reported putamen-*right* middle frontal gyrus connectivity), and a decrease in dACC-precuneus connectivity to a different portion of the precuneus than previously reported. We interpret this as a partial replication of these earlier reports, especially in light of important methodological differences between our study and the previous ones.

Two studies (Nasseri et al., 2020; Sharma et al., 2020) included women using a variety of OCPs, which may have included important differences in their formulation (e.g., different levels of androgenicity), as compared to our study in which all participants used the same formulation. Additionally, while the OCP phase was uniform across all studies, the comparator group phase was not. Nasseri et al. used the inactive pill phase as a comparator group, and Sharma et al. included naturally-cycling women in several menstrual phases. Importantly, Nasseri et al. measured resting-state networks following an acute stressor, which our study did not include. Our paradigm was most similar to Engman et al., and used the same formulation of OCPs, but differed in the study design (cross-sectional vs. longitudinal). We also elected to use Schaefer nodes for the selection of seed ROIs, which overlapped with but were not identical to the seed selection method in these other studies. These methodological differences may explain part or all of the discrepancies in findings.

The findings of our amygdala seed-based analysis are overall inconsistent with the results of Engman et al. (2018). Potential functional connectivity changes involving the amygdala are nonetheless hard to ignore given the amygdala’s role in fearful emotions and anxiety disorders (Hu et al., 2020; Phelps & LeDoux, 2005). Because of the higher propensity for women to experience changes in mood symptoms, which can include anxiety and anxiety-related symptoms during the luteal phase of the menstrual cycle and sometimes in response to hormonal contraceptives (Handy et al., 2022; Mu & Kulkarni, 2022; Reynolds et al., 2018), amygdala activity is a prime candidate as the potential cause of negative emotions in response to OCPs. Interestingly, a recent study was similarly unable to replicate the decrease in amygdala-postcentral gyrus connectivity, finding such a decrease originating from the dACC seed instead (Hidalgo-Lopez et al., 2023).

Given the partial or non-replication of previously reported seed-based effects, we turned to an exploratory, data-driven approach to investigate other potential effects of OCPs on functional connectivity.

#### 4.2.1. Whole-Brain Connectome Analysis

In addition to the a priori replication efforts described above, to fully leverage this unique data set, we also performed a series of hypothesis-generating exploratory analyses. To balance the risks of type 1 and type 2 errors, we applied a series of dimension reduction approaches. First, we provide a visual overview of the effect size of edge changes, both with respect to each individual node, and also averaged across the nodes within previously-defined canonical networks. These show that, at a network level, OCPs produce the largest increases in connectivity within the dorsal attention network, and the largest decreases in connectivity between the somatomotor network and visual network, with a comparable decrease in connectivity between the somatomotor network and default mode network. However, inferential statistics were not performed and therefore these should be considered descriptive rather than hypothesis-confirming findings.

#### 4.2.2 Functional Connectome Fingerprint Identifiability

Functional connectome fingerprint results showed an increase in *I_Other_*during OCP usage while *I_Self_* and *I_Diff_* remained statistically similar. An increase in similarity between subjects without altering self-identifiability suggests that, although idiosyncratic edges remain respective to their participants, OCPs cause a convergence towards a common effect on FC. This is consistent with previous findings from a single-subject longitudinal study, which also identified an overall effect of OCPs on network connectivity dynamics (Jensen et al., 2022). Interestingly, the result from Jensen et al. showed a generalized increase in edge connectivity in the brain during OCP use with less organized network segmentation, which is consistent with our finding that ICC values generally fell brainwide.

Additionally, although a general trend of self-identifiability is still observed for each subject across the different interventions, the identification is much less robust relative to within-scan comparisons, as shown by Figure 5. Furthermore, the assessment of *I_Self_*, *I_Diff_*, and *I_Other_* using the progressive inclusion of the top ICC-value edges from the placebo group shows larger differences between interventions in all three metrics on smaller inclusion scales. These differences highlight the effects of OCPs on idiosyncrasy, since common and presumably unrelated edges are effectively ignored.

The breadth of the network-wise ICC changes make specific, localized conclusions difficult to determine. However, these changes themselves show how an ROI-based method of analyzing OCPs may not be sufficient to study their varying effects on users. Our fingerprint results give support to the narrative that OCPs have a generalized effect on the whole brain, and the changes in the idiosyncratic characteristics of the users themselves may explain more variance in subjective experience than any single ROI-ROI connection can. As the neuroimaging field broadly makes an effort to move toward precision medicine in neuroscience and psychiatry, identifying these idiosyncratic patterns could inform more individualized treatment plans.

Connectome fingerprinting has proven effective in studying the effects of pharmaceuticals with widespread connectivity changes (Tolle et al., 2024), and our results indicate OCPs belong to this category. Neuroscience and psychiatry alike are transitioning towards precision medicine to treat individuals based on their own specific characteristics, recognizing the complexity of a dynamic system like the brain (Hampel et al., 2023). Such medicine could greatly benefit from a database of fingerprint profiles of both drugs and patients. Therefore, profiling the effects of OCPs on idiosyncrasy could lead to more precise and tolerable prescriptions issued to each user, helping the problem of unpredictable reactions experienced by users.

#### 4.2.3. Brain-behavior relationships: Network changes underlying changes in mood

We observed changes in connectivity that correlated with changes in mood symptoms in nine nodes that are distributed throughout the brain. Some nodes have plausible pre-existing relationships with networks or emotional symptoms. The right posterior cingulate cortex is a key node of the default mode network, therefore positioned to exert control over resting emotional states and self-perception (Fransson & Marrelec, 2008; Maddock et al., 2003). It is also implicated in a number of neuropsychiatric and degenerative diseases (Greicius, 2008; Leech & Sharp, 2014), making it a logical target for future analysis of rsFC during OCP-related negative behavioral symptoms. We also observed connectivity-symptom correlations in regions of the prefrontal cortex. The dorsolateral prefrontal cortex has been shown to have different activity between women with and without PMDD (Gingnell et al., 2013; Petersen et al., 2018), with additional evidence implicating its importance in regulating resting state mood, perception of emotion during use of working memory, anxiety symptoms, and other DRSP measures (White et al., 2023). This and other, nearby prefrontal regions could be involved in a pathway of emotional dysfunction, causing certain users of OCPs to experience negative symptoms. We believe these data suggest that more work focusing on prefrontal contributions to emotional problems while using OCPs is warranted.

We also observed parietal effects involving the precentral gyrus and precuneus. The inferior parietal lobe has been robustly linked to menstrual cycle influences on brain structure and functional connectivity (Dubol et al., 2021). The precentral gyrus is more typically associated with motor function (Hartwigsen et al., 2012; Hartwigsen & Siebner, 2015), while the precuneus has a more broadly-accepted role in neuropsychiatric symptoms (Dadario & Sughrue, 2023). Together, these data point toward an effect of OCPs on a broad, distributed network that influences regions linked to emotional regulation and sensorimotor integration. Therefore, it may be worth including regions related to emotions in the investigation of negative symptoms related to OCP use. Future studies may be able to reveal a more specific, hormone-sensitive brain network that governs negative emotional responses to OCPs. We propose that such a hormone-sensitive network could link negative affective responses to hormone signaling transdiagnostically – i.e., the same network could be implicated in negative emotional responses to OCPs, to hormonal fluctuations during the menstrual cycle, to postpartum hormone changes, and other experiences or exacerbations of negative affect that take place during hormonal transitions. We emphasize that this is a hypothesis for future study rather than a conclusion to be drawn from these data.

### 4.3 Limitations

This study has several limitations, starting with the small sample size (*N* = 26), which limits generalizability of the findings and raises the possibility of both type I and type II errors. Second, the study focused on only one oral contraceptive formulation (30 μg ethinyl estradiol/0.15 mg levonorgestrel), which may limit the applicability of these findings to other types of oral contraceptives with different hormone compositions. Brain connectivity changes were observed at rest, but measuring brain function during an emotional task might have provided more robust or meaningful observations, potentially linking neural changes more directly to emotional or cognitive outcomes. Additionally, the exploratory analyses suggest potential neural correlates of oral contraceptive use, but definitive conclusions cannot be made without further replication in larger, independent cohorts. The relatively short intervention period (21 days) may not capture long-term effects of oral contraceptive use on brain networks, which could differ with extended exposure to the hormones. Finally, OCPs have cardiovascular effects, and we cannot rule out the possibility that the observed effects on functional connectivity are secondary to cerebrovascular effects of OCPs

### 4.4. Implications

Our findings carry potential implications for both clinical practice and future research. Clinically, identifying the brain networks most affected by OCPs could lead to more personalized contraceptive prescriptions, minimizing mood-related side effects, improving patient satisfaction, and reducing the likelihood of unwanted pregnancies. Given the wide variety of OCP formulations, the ability to prospectively stratify patients to the options least likely to cause unwanted side effects would be highly beneficial. We propose that achieving this goal will likely require a multivariate method, such as functional connectome fingerprinting. For researchers, considering the widespread use of OCPs and their apparent substantial influence on the connectome, researchers may wish to account for this source of variance to optimize the design and interpretation of their neuroimaging investigations.

Finally, while we observe widespread influences of OCPs on the connectome, we also identified a smaller set of brain regions that appear to link connectivity changes with mood alterations. We propose that these regions may be part of a hormone-sensitive network, potentially acting as a transdiagnostic mechanism underlying negative emotional responses across various conditions. We encourage future studies to verify or reject the brain regions implicated in this investigation.

### 4.3 Conclusion

In conclusion, this study sought to replicate previously reported effects of oral contraceptive pills (OCPs) on brain connectivity, yielding mixed but predominantly null findings. Although key connectivity changes in regions such as the amygdala and ventromedial prefrontal cortex were not replicated, the observed changes in connectivity between the putamen and the middle frontal gyrus may offer partial support for earlier studies. The data-driven exploration identified several additional brain regions where OCPs appear to influence functional connectivity, with potential implications for understanding the neural basis of mood changes associated with hormonal contraceptive use. Notably, OCPs were found to reduce individual variability in functional connectivity, suggesting a converging effect on brain network organization, and demonstrating the utility of functional connectome fingerprinting as a robust method to measure within-subject changes in the functional connectome. Together, these network-based results suggest that OCPs exert a widespread effect on functional connectome organization, rather than concentrating effects on a single or handful of ROIs.

The constellation of findings presented here further demonstrates the critical need for larger sample sizes to draw more definitive conclusions. This highlights the importance of data sharing, collaborative consortia studies, and greater investment from funding agencies to support large, adequately powered studies.

## Data and Code availability

The code used in these analyses is available on Github (https://github.com/HumanBrainZappingatUCLA/OCPRSNs). Any additional information required to reanalyze the data used in this paper is available upon request and use agreement with the corresponding author.

## Author Contributions

Conceptualization, T.J. and N.P.; Methodology, G.H. and T.J.; Software, J.L. and T.J.; Formal Analysis, G.H., J.L., N.P., and T.J.; Investigation, N.P.; Data Curation,G.H., J.L. and T.J.; Writing – Original Draft, G.H., T.J. and N.P.; Writing – Review & Editing, G.H., T.J., E.D.L., A.R., and N.P.; Visualization, G.H. and T.J.; Supervision, T.J. and N.P.; Project Administration N.P..

## Funding

This study was supported by a grant from the National Institutes of Health (NIDA), R21DA040168 to EDL.

## Declaration of Competing Interests

The authors declare no competing interests.

## Acknowledgements

We thank the staff of the Center for Cognitive Neuroscience for providing aid and support for all fMRI imaging sessions.

## Notes

### Competing Interest Statement

The authors have declared no competing interest.

### Summary of Updates

This version of the manuscript has been revised to update the following: Statement of IRB approval has been added.

