## Supplement for "Effects of oral contraceptive pills on brain networks: A replication and extension"

### Supplemental Materials

*1. Replication efforts – connectivity of* a priori *nodes, additional detail:*

In addition to evaluating seed-based connectivity of *a priori* nodes by matching regions to the corresponding Schaefer atlas nodes, we used a second strategy to test the hypotheses using an alternative ROI definition. In this alternative approach, we used average region-of-interest (ROI) masks from Neurosynth.org (Neurosynth.org), which were thresholded and binarized using FSL to distinguish the ROIs with lateral specificity. The left amygdala mask was thresholded to a z-score value of 8.5 and adjusted to include the values on the left side only. The right putamen mask was thresholded at 8.7 and similarly segregated to voxels in the right putamen. The left parahippocampal mask was thresholded to 9 and included only the left parahippocampal gyrus. The dACC mask on Neurosynth.org contained sparse voxels far outside of the desired ROI after thresholding, therefore we used a binarized map of the ACC overlaid by the thresholded dACC to remove outlier points and obtain a functional map exclusive to voxels in the dACC. These node definitions are shown below in Supplemental Figure S1. This approach did not change the results, and we did not replicate previous findings using this ROI definition strategy.

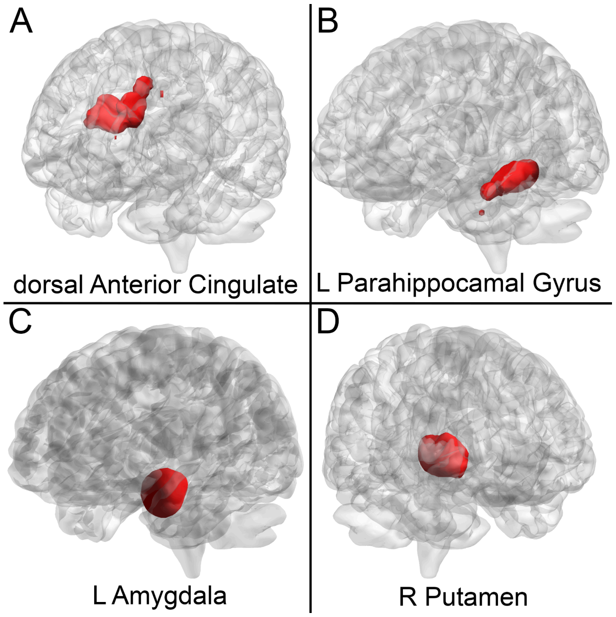


**Supplemental Figure S1:** A priori nodes using functionally-defined regions derived from Neurosynth functional connectivity maps.

*Additional findings:*

a. *Right putamen connectivity:* In a previous analysis, Sharma et al. reported higher connectivity between the right putamen and right middle frontal gyrus. We were not able to replicate this effect – however, we observed an effect in the contralateral middle frontal gyrus. The MNI coordinate of the previous report (MNI: 30, 12, 60) and the Schaefer node in which we observed this effect are visualized below in Supplemental Figure S1.

***Supplemental Figure S2:*** Panel A shows the MNI coordinate 30, 12, 60 (previously reported effect). Panel B shows Schaefer node 186 (currently observed effect).

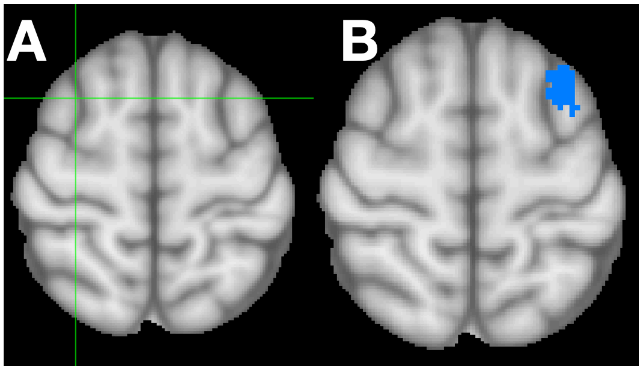


b. *Left dorsal anterior cingulate connectivity:* In a previous analysis, Engman et al. reported lower dACC-precuneus connectivity in women using OCPs compared to placebo (during the luteal phase). We did not replicate this finding, but did show lower connectivity dACC-during OCPs vs. placebo in a nearby portion of the precuneus.

***Supplemental Figure S3:*** Panel A shows the MNI coordinate 0, -54, 63 (location of previously reported dACC-precuneus connectivity change). Panel B shows Schaefer node 195 (location of currently observed dACC-precuneus connectivity change).

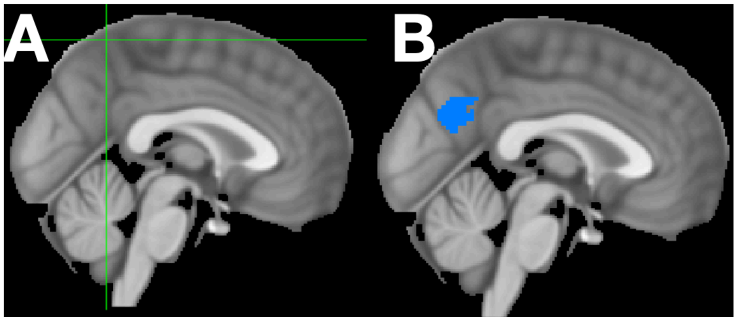


*2. Attempts at data reduction – exploring connectivity changes resulting from OCP use:*

Considering connectivity changes between each node in the Schaefer parcellation produces 175,561 (419 x 419) comparisons. Because of the nature of brain networks, many of these are likely to be collinear/non-independent. Therefore, we first wanted to test our assumption that OCPs have *any* effect on functional connectivity, before exploring *which* functional connectivity effects could be plausibly attributed to OCPs.

If the effect of OCPs on functional connectivity was null, then the overall distribution of effect sizes (for the effect of OCPs on change in each edge) would be expected to fit an approximately Gaussian distribution. We plotted a histogram of effect sizes (all nodes) and observed a leftward shift with heavy tails. We applied the Kolmogorov-Smirnov test to compare the observed distribution to a normal distribution and found that the two differed significantly, p = 2.2391e-37. We also plotted the distribution of observed effect sizes against the normal distribution in a Q-Q plot. The observed (sample) and theoretical distributions again differ, as can be seen visually by the non-overlapping lines. Together, these data indicate that OCPs do indeed alter functional connectivity (code currently: OCP_RSN_distribution.ipynb in my home directory).

***Supplemental Figure S4:*** The first panel shows the overall distribution of effect sizes (effect of OCPs on each edge). The distribution peaks slightly to the left of zero, and differs significantly from a normal distribution. The Q-Q plot to the right visualizes the same information differently, showing substantial deviations from the normal distribution in both the right and left tails, where larger effect sizes are represented. This suggests that OCPs produce more effects in each tail – i..e, both increases and decreases in connectivity – than would be expected by chance.

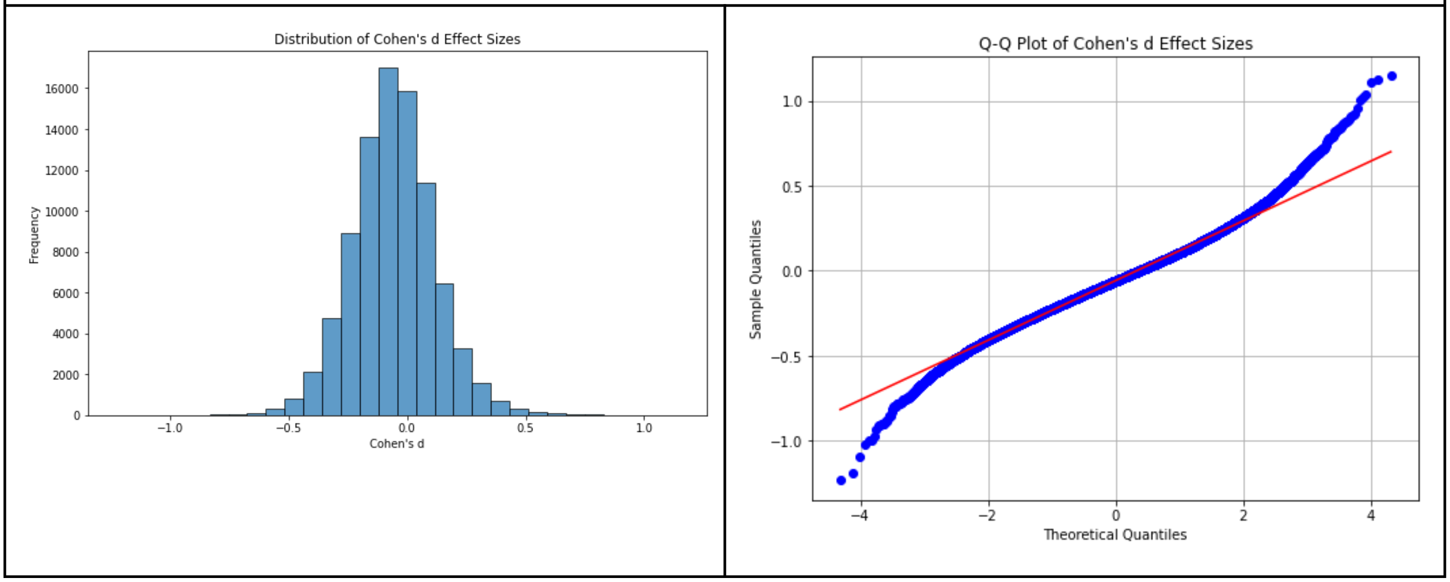


Because the time series from some nodes are very likely to be collinear with one another, we subsequently attempted dimensionality reduction using principal component analysis. First, we calculated connectivity matrices and the change therein as previously described in the manuscript. Then, we used the PCA implementation in the python library *scikit-learn*. To identify the optimal number of components for analysis, we calculated the cumulative variance explained by each additional component, then identified the inflection point of this curve by taking the second derivative of the curve's first derivative. This showed a clear inflection at 26 components (which likely corresponds to the 26 participants entered into the PCA). The majority of the variance (67.25%) was explained by the first component. However, the feature loadings for PC1 were approximately evenly distributed, suggesting that the connectivity changes observed were driven by many small-moderate changes rather than isolated, large changes (code available OCs_PCA_exploratory.ipynb in my home directory; move to GitHub before manuscript release).

**Supplemental Figure S5:** The same data shown in manuscript Figure 10 are shown in separate plots here. Approximate node identities are given in the table below. Some nodes span more than one cortical region (using other definitions of regions). Exact node geographies and positions can be seen by accessing the Schaefer 400 node parcellation.

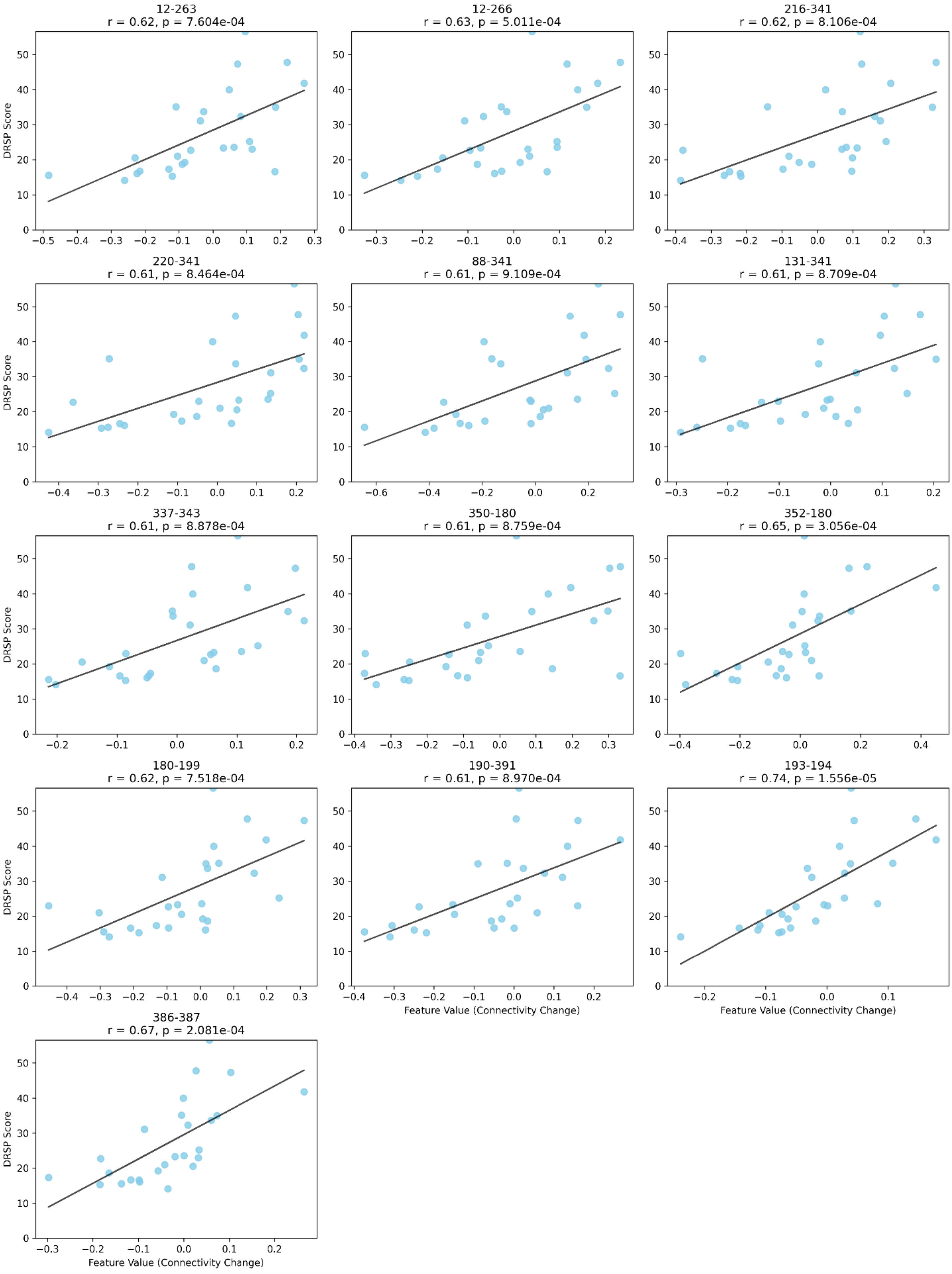
12-263
r = 0.62, p = 7.604e-04
12-266
r = 0.63, p = 5.011e-04
216-341
r = 0.62, p = 8.106e-04
220-341
r = 0.61, p = 8.464e-04
88-341
r = 0.61, p = 9.109e-04
131-341
r = 0.61, p = 8.709e-04
337-343
r = 0.61, p = 8.878e-04
350-180
r = 0.61, p = 8.759e-04
352-180
r = 0.65, p = 3.056e-04
180-199
r = 0.62, p = 7.518e-04
190-391
r = 0.61, p = 8.970e-04
193-194
r = 0.74, p = 1.556e-05
386-387
r = 0.67, p = 2.081e-04
DRSP Score
Feature Value (Connectivity Change)

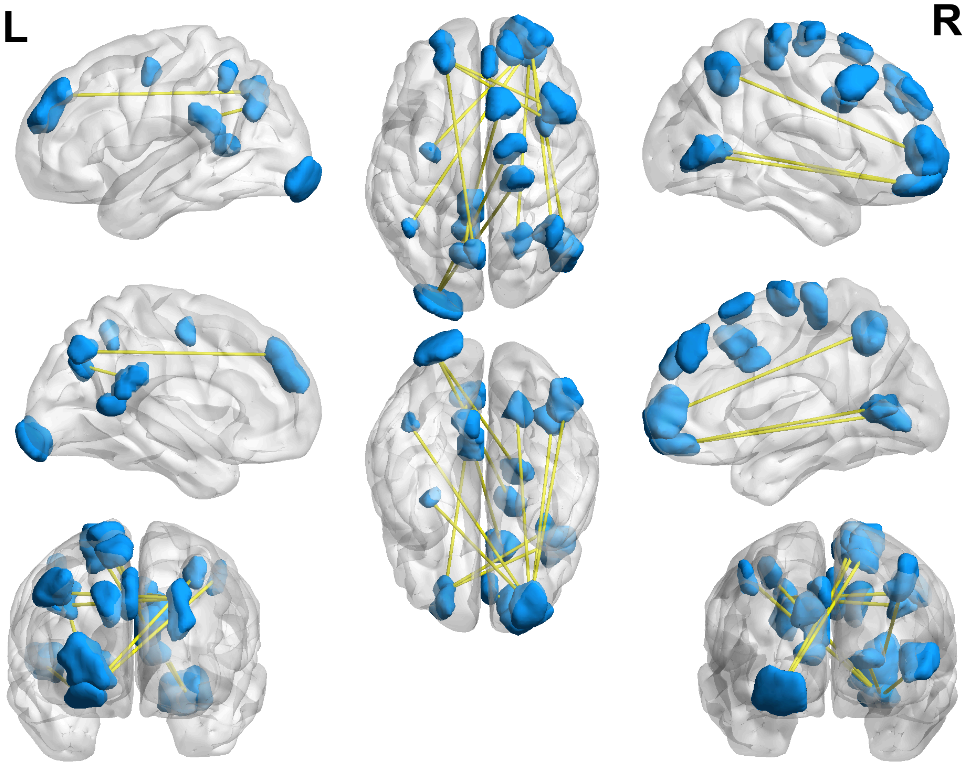


**Supplemental Figure S6: Nodes involved in connectivity changes that correspond to DRSP changes.** The nodes span prefrontal, parietal, and occipital cortices. Moving from anterior to posterior, prefrontal cortex nodes include the frontal pole, superior frontal gyrus, middle frontal gyrus, and precentral cortex. Parietal regions include posterior cingulate cortex, the angular gyrus, and precuneus. Occipital regions include the lingual gyrus and visual cortex (approximately V5).

| **Node legend** | |
|---|---|
| 12 | Occipital pole |
| 88 | Left precentral gyrus |
| 131 | Left angular gyrus |
| 180 | Left frontal pole |
| 190 | Left posterior cingulate |
| 193 | Left posterior cingulate |
| 194 | Left precuneus |
| 199 | Left precuneus |

| Node legend | |
|---|---|
| 216 | Right lateral occipital cortex |
| 220 | Right lingual gyrus |
| 263 | Right precentral gyrus |
| 266 | Right superior frontal gyrus |
| 337 | Right superior occipital cortex |
| 341 | Right frontal pole |
| 343 | Right frontal pole |
| 350 | Right middle frontal gyrus |
| 352 | Right middle frontal gyrus |
| 386 | Right frontal pole |
| 387 | Right superior frontal gyrus |
| 391 | Right superior frontal gyrus |
